# NFATc3 Enhances DREAM Complex-Driven Transactivation

**DOI:** 10.64898/2026.01.12.699082

**Authors:** Jieun Ahn, Yoojeong Seo, Jinho Jang, Bongjun Kim, Moonjong Kim, Jie Zhang, Sohee Jun, Jae-Il Park

## Abstract

The nuclear factor of activated T cells (NFAT) transcription factors, activated by calcium-calcineurin signaling, regulate various cellular processes, including cell differentiation, angiogenesis, and immune cell activation. Nonetheless, their roles in tumor cells remain largely undefined. The dimerization partner, RB-like, E2F, and multi-vulval class B (DREAM) complex orchestrates cell quiescence and proliferation. Pharmacological mimicry of DREAM-activated transcriptional signatures identified two calcium signaling inhibitors that suppressed lung adenocarcinoma (LUAD) cell proliferation. Among the five NFAT members, NFATc3/NFAT4 was predominantly expressed in LUAD cells and required for both LUAD cell proliferation and DREAM target gene transactivation. NFATc3 was enriched at DREAM target promoters and associated with the DREAM complex, possibly via LIN9, a scaffolding protein of the Multi-Vulva class B (MuvB) core proteins. These findings reveal an unexpected role for NFATc3 in promoting DREAM target gene transactivation and suggest the calcium-NFATc3 axis as a molecular target in LUAD, enriched by DREAM complex activation.

## Introduction

Lung adenocarcinoma (LUAD), the most prevalent subtype of non-small cell lung cancer, accounts for approximately 40% of all lung cancer cases^1^. Mutations in *KRAS* (∼33%) and *TP53* (∼55%) are frequently observed in LUAD^2^. These genetic mutations are sufficient to induce lung tumors in mice^3, 4^. Although targeting RAS signaling or restoring TP53 function has been explored for cancer treatment, such clinical applications remain challenging^5, 6^. Despite recent advances in lung cancer research, therapeutic options are still limited, and patient survival rates have not significantly improved. Therefore, developing new and viable treatments for lung cancer is imperative.

Somatic cells remain quiescent upon terminal differentiation, and disruption of this process leads to exit from cell quiescence, a condition strongly implicated in cancer^7, 8^. The dimerization partner, RB-like, E2F, and multi-vulval class B (DREAM) complex (also called the DRM complex in *Caenorhabditis elegans* and the dREAM complex in *Drosophila melanogaster*) is an evolutionarily conserved multiprotein complex that orchestrates cell quiescence and the cell cycle^9–12^. In association with RBL2/p130 (retinoblastoma-like protein 2), E2F4, and DP1 (E2F dimerization partner 1), the DREAM complex localizes to the promoters of cell cycle-regulating genes, repressing their transcription and inducing quiescence via G0 and G1 arrest^12^. Upon dissociation of RBL2, E2F4, and DP1 from the DREAM complex, the remaining multi-vulval class B (MuvB) core complex (composed of LIN9, LIN37, LIN52, LIN54, and RBBP4) binds to BMYB and FOXM1, which transactivates cell cycle-related genes at the S/G2/M phases^13^. Given its crucial role in cell cycle regulation, dysregulation of the DREAM complex is associated with various cancers^14–16^. Consistently, the overexpression of DREAM complex targets, including FOXM1, BMYB, PLK1, TOP2A, and CCNB1, is frequently observed in many cancers. However, the mechanism behind the aberrant regulation of the repressive DREAM complex in cancer cells remained elusive until our recent study^17^. Proliferating cell nuclear antigen (PCNA)-associated factor (PCLAF; also known as PAF/KIAA0101) is a small nuclear protein consisting of 111 amino acids^18^. Through its interaction with PCNA, PCLAF regulates DNA replication and repair^19, 20^. Studies, including ours, have shown that PCLAF is highly upregulated in many cancers but scarcely expressed in normal cells^18, 21–29^. PCLAF expression is essential for cancer cell proliferation^19, 22, 23, 25, 28, 29^ and cell stemness^26, 30, 31^. Recently, we uncovered the in vivo role of PCLAF in lung cancer. The PCLAF-remodeled DREAM complex is crucial for KRAS- and TP53-driven lung tumorigenesis^17^. Another study showed that Chloride Intracellular Channel 4 (CLIC4), a DREAM target gene and an enhancer of TGF-β signaling, drives alveolar cell plasticity for lung regeneration^32^, identifying an unexpected role for the DREAM complex in modulating tissue regeneration.

To identify small molecules that recapitulate the PCLAF-depleted transcriptome and phenotype, we exploited Connectivity Map (CMAP) approaches^33^ with RNA-seq datasets from mouse and human lung cancer cells. Interestingly, we discovered that Ca^2+^ signaling inhibitors, cyclosporin A (CsA) and NNC 55-0396, significantly inhibited DREAM target gene expression and cell growth. Intriguingly, these results are consistent with our previous findings where CsA, acting as a pharmacological inhibitor of the PCLAF-DREAM axis, suppressed lung tumorigenesis *in vivo* by inducing p130-dependent G0/G1 arrest^17^.

Intracellular Ca^2+^ regulates various cellular processes, including the cell cycle, transcription, exocytosis, motility, and apoptosis^34^. NNC 55-0396 is an inhibitor of the T-type Ca^2+^ channel^35^ and has been shown to inhibit cancer cell growth and angiogenesis^36–38^. However, its impact on lung cancer remains unknown. CsA binds to cyclophilin and inhibits calcineurin to block Ca^2+^ signaling^39^. Both CsA and NNC 55-0396 effectively induced G0/G1 cell cycle arrest and downregulated DREAM target gene expression in LUAD cells^17^. These findings suggest that blocking Ca^2+^ signaling may suppress the PCLAF-DREAM axis, thereby inhibiting lung cancer cell growth.

The activation of Ca^2+^ signaling is strongly implicated in cancer^40–42^. However, its oncogenic roles in lung tumorigenesis remain unclear. Increased intracellular Ca^2+^ activates calcineurin, a Ser/Thr phosphatase, which dephosphorylates NFATs, exposing their nuclear localization signals. This allows NFATs to translocate into the nucleus and activate downstream target genes^43^.

In this study, we determined the impact of the calcium pathway in lung tumor cell proliferation and further demonstrated that NFATc3 physically and functionally interacts with the DREAM complex, promoting the transactivation of DREAM target genes for LUAD cell proliferation.

## Results

### Calcium signaling is required for LUAD cell proliferation

Elevated calcium levels are frequently observed in various cancers^44^. We previously demonstrated that Cyclosporine A (CsA), a calcineurin inhibitor, exerts a growth-inhibitory effect on LUAD cells^17^. To explore therapeutic candidates based on these findings, we leveraged our prior CMAP analysis, which integrated transcriptomic profiles from human H1792 and mouse KP (*Kras*^G12D^; *Trp53*^Δ/Δ^) LUAD cells, identifying compounds that possibly reverse the LUAD oncogenic signature (Fig. 1A). Among the high-ranking candidates, we specifically focused on compounds associated with calcium signaling pathways: CsA and BAPTA-AM, a cell-permeable intracellular Ca^2+^ chelator (Fig. 1B). To validate their effects, these two candidates were tested with FK506 (Tacrolimus), another well-established calcium signaling inhibitor, on LUAD cell lines. H1792 and A549 LUAD cells treated with CsA, FK506, or BAPTA-AM exhibited growth arrest in the G0/G1 phase (Fig. 1C, D). Furthermore, we found that these inhibitors reduced NFATc3 expression, a key Ca^2+^ signaling mediator, whereas the KRAS-G12C inhibitor AMG510 had no effect on NFATc3 levels (Fig. 1E). These findings suggest that Ca^2+^ signaling promotes LUAD cell growth.

**Figure 1.**
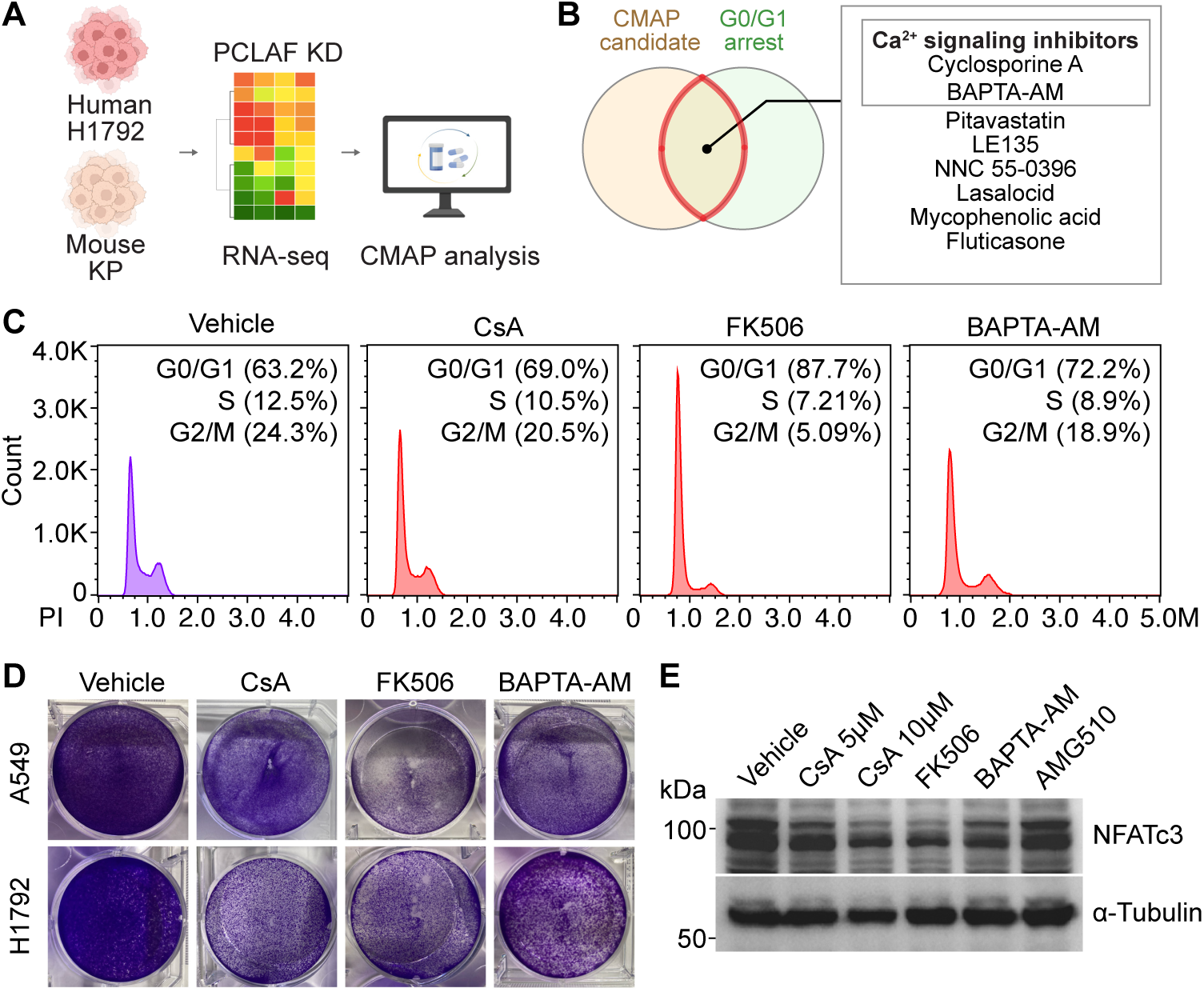
LUAD growth inhibition by Ca^2+^ signaling inhibitors. **A.** Schematic workflow of the CMAP-based drug discovery process. **B.** Venn diagram showing the overlap between candidate compounds identified via CMAP and associated with growth arrest. **C, D.** H1792 cells were treated with each inhibitor (24 hrs) and analyzed by flow cytometry (C) and crystal violet staining (D). Representative images (n ≥ 3) are shown. **E.** Immunoblot (IB) analysis of A549 cells treated with CsA, FK506, BAPTA-AM, or AMG510. AMG510 served as a negative control.

### NFATc3 mediates calcium signaling-dependent LUAD cell proliferation

We previously demonstrated that CsA-mediated suppression of the DREAM complex inhibits lung tumorigenesis^17^. To further validate these pharmacological findings using genetic models and to identify the specific molecular mediators involved, we focused on the NFAT family, which are key downstream effectors of Ca^2+^-calcineurin signaling^43^. Among the five NFAT family members (NFATc1, NFATc2, NFATc3, NFATc4, and NFAT5), *NFATc3* expression was found to be relatively enriched in human LUAD cells (Fig. 2A), human LUAD tissues (Fig. 2B), and murine LUAD samples (Fig. 2C), which led us to determine whether NFATc3 is essential for LUAD cell proliferation. We depleted endogenous NFATc3 using lentiviruses encoding shRNAs targeting NFATc3 (Fig. 2D). Compared to the control (shCtrl), NFATc3-depleted LUAD cells (A549 and H1792) exhibited significant growth inhibition (Fig. 2E), as confirmed by flow cytometry analysis (Fig. 2F). These findings suggest that the calcium-NFAT signaling axis is crucial for LUAD cell growth.

**Figure 2.**
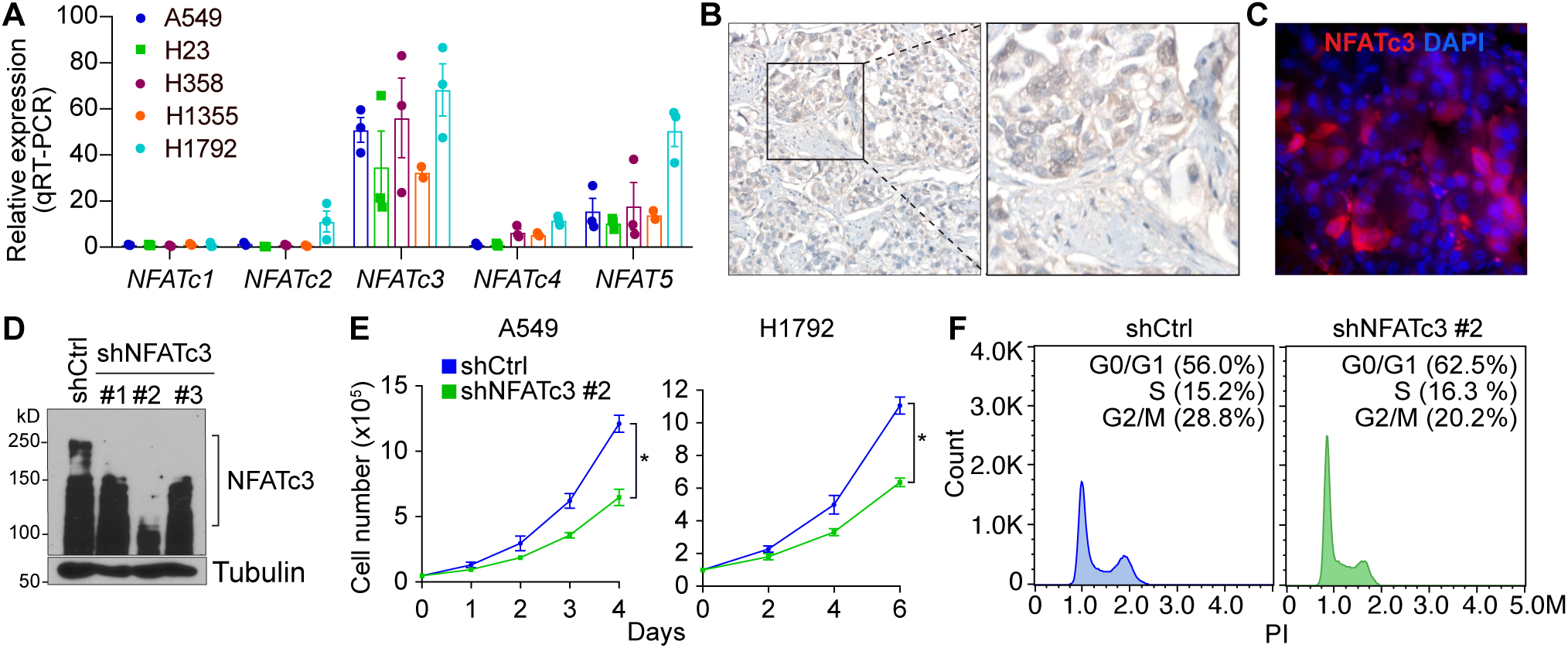
NFATc3 is required for LUAD cell proliferation. **A.** qRT-PCR of five LUAD cells for NFAT family genes. **B.** Immunostaining of human LUAD tissue microarray for NFATc3. **C.** staining of murine LUAD isolated from *Kras^G12D^ Trp53*^-/-^ mice. **D.** Immunoblot (IB) of H1792 cells stably transduced with lentiviruses encoding shCtrl (control) or shRNAs against NFATc3. **E.** A459 and H1792 cells (shCtrl vs. shNFATc3) were examined for cell proliferation by cell counting. **F.** Flow cytometry analysis. Representative images (n ≥ 3) are shown. *P* values were calculated using Student’s *t*-test; error bars, standard deviation (SD).

### NFATc3 occupies and activates the DREAM complex target promoters

We previously reported that PCLAF remodels the DREAM complex to drive LUAD cell hyperproliferation^17^. Notably, our findings showed that calcium signaling inhibitors, CsA and NNC 55-0396, significantly inhibit DREAM target gene expression^17^, and various calcium inhibitors were further confirmed to suppress LUAD cell proliferation (Fig. 1). Given the requirement of NFATc3, a calcium-activated transcription factor, in LUAD cell proliferation (Fig. 2), we hypothesized that NFATc3 may positively regulate DREAM target genes. To test this, we depleted NFATc3 in H1792 cells and assessed the expression of DREAM target genes. NFATc3 knockdown led to a transcriptional downregulation of the DREAM target genes (Fig. 3A). Cleavage Under Targets & Release Using Nuclease (CUT&RUN)-qPCR assays revealed NFATc3 enrichment on the promoters of DREAM target gene including *TOP2A*, *NEK2*, and *UBE2C* (Fig. 3B). Furthermore, Chromatin immunoprecipitation (ChIP)-sequencing analysis of the publicly available datasets indicated the co-enrichment of LIN9, a core component of the DREAM complex, and NFATc3 on DREAM target gene promoters (*Kif22* and *Ncapd2*) (Fig. 3C). These findings suggest that NFATc3 is functionally involved in the transactivation of DREAM target gene promoters.

**Figure 3.**
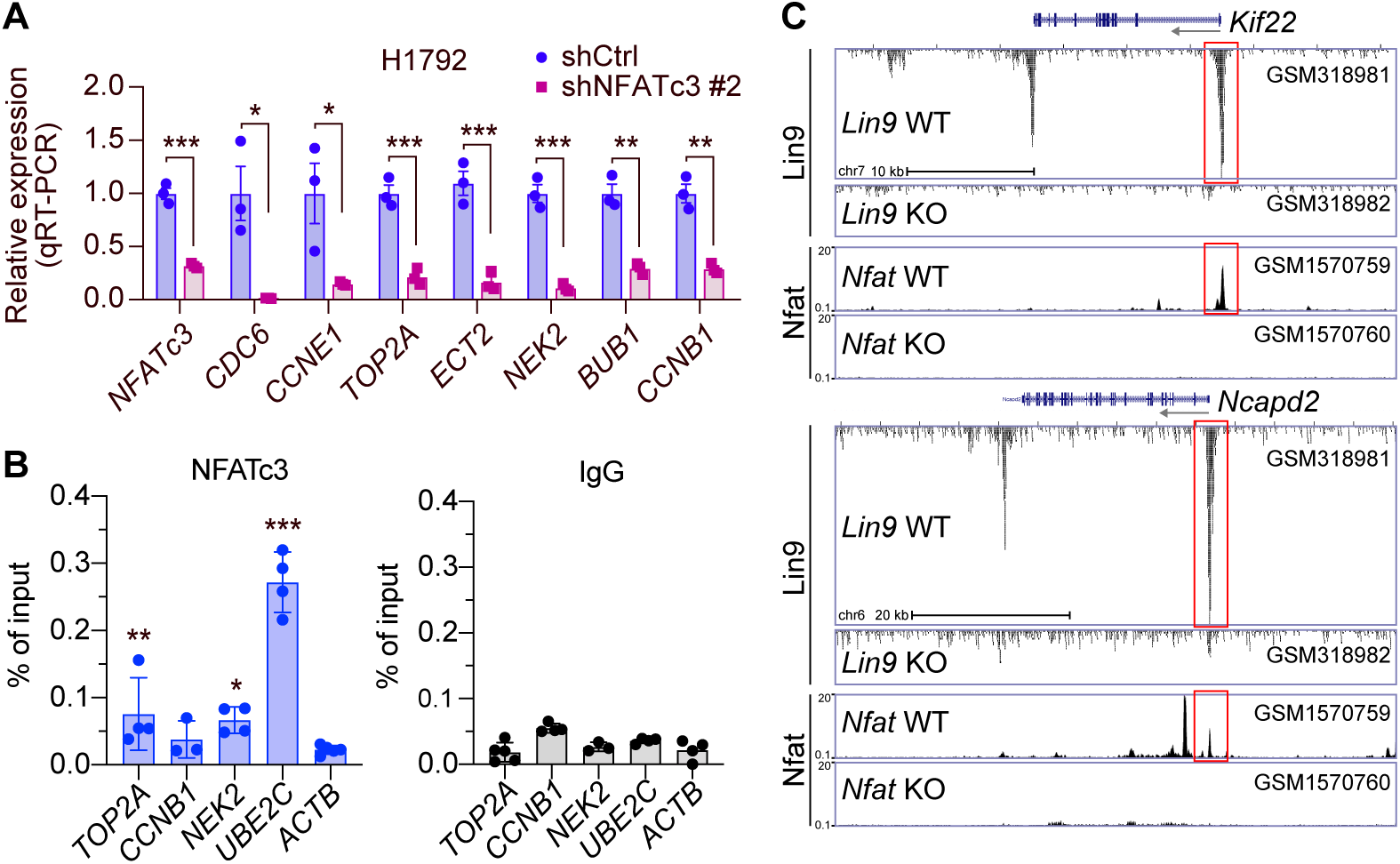
DREAM target gene promoter activation by NFATc3. **A.** qRT-PCR of H1792 cells stably transduced with shCtrl (control) or shNFATc3 #2 (clone number) for *NFATc3* and the DREAM target genes. **B.** CUT&RUN assays of A549 cells with anti-NFATc3 antibodies or IgG (control). The *ACTB1* promoter served as a negative control. **C.** Promoter enrichment of transcription factors (TFs) on the DREAM target genes (*Kif22* and *Ncapd2*). Each dataset was visualized on the UCSC genome browser. The dataset accession numbers were shown. Red boxes indicate the co-enrichment of Lin9 and Nfat on the promoters. *P* values were calculated using Student’s *t*-test; error bars, standard deviation (SD).

### NFATc3 binds to the DREAM complex

Having identified the enrichment of NFATc3 on DREAM target gene promoters (Fig. 3), we next investigated whether NFATc3 physically interacts with the DREAM complex. AlphaFold-multimer modeling predicted that the MuvB core serves as a structural scaffold for NFATc3 integration (Fig. 4A, B). Specifically, the predicted aligned error (PAE) plot indicated a high-confidence interaction between NFATc3 C-terminal (401-700 amino acid [AA]) and the N-terminal region of LIN9 (120-268 AA) and LIN37. LIN9 (120-268 AA) serves as a docking site for RBBP4 and LIN37, while the LIN9 (353-434 AA) and LIN9 (443-545 AA) regions anchor LIN52 and LIN37-LIN54, respectively. This prediction aligned with established models^45^, confirming that the LIN9 module provides an essential platform for recruitment of regulatory proteins (Fig. 4C). Our protein-protein interaction (PPI) analysis further confirmed the specific domain requirements for NFATc3-MuvB complex (Fig. 4D, E). NFATc3 (401–700 AA) anchors to the internal domain of LIN9 (120-268) with high confidence. The interaction of MuvB-associated transcriptional activators, FOXM1 and MYBL2, with NFATc3-MuvB complex was also predicted by AlphaFold (Fig. 4F, G). Notably, interface analysis revealed that the amino acid K522 of NFATc3 was in close proximity to FOXM1 V233 and H234, and these residues were highly conserved across vertebrates, suggesting a functionally critical contact point for assembly (Fig. 4G, H). To validate this in silico prediction, we performed endogenous co-immunoprecipitation (co-IP) in A549 cells. NFATc3 interacted with key DREAM components, including LIN37, LIN9, RBBP4, FOXM1, and LIN54 (Fig. 4I, Supplementary Fig. 1A, B). To evaluate the co-regulatory potential of NFATC3 and FOXM1, we performed a motif scanning analysis on the promoter regions (-500 to +100 bp) of 9 DREAM target genes using the JASPAR database and FIMO (Find Individual Motif Occurrences) tool. The integrated bubble plot reveals a high density of binding clusters for both transcription factors, particularly concentrated within the -400 to -200 bp upstream region relative to the TSS (Fig. 4J). The spatial proximity of NFATC3 and FOXM1 binding sites in several promoters suggests a potential synergistic transcriptional regulation. These findings suggest that NFATc3 binds to core components of the DREAM/MuvB complex to drive the transactivation of DREAM target genes (Fig. 4K).

**Figure 4.**
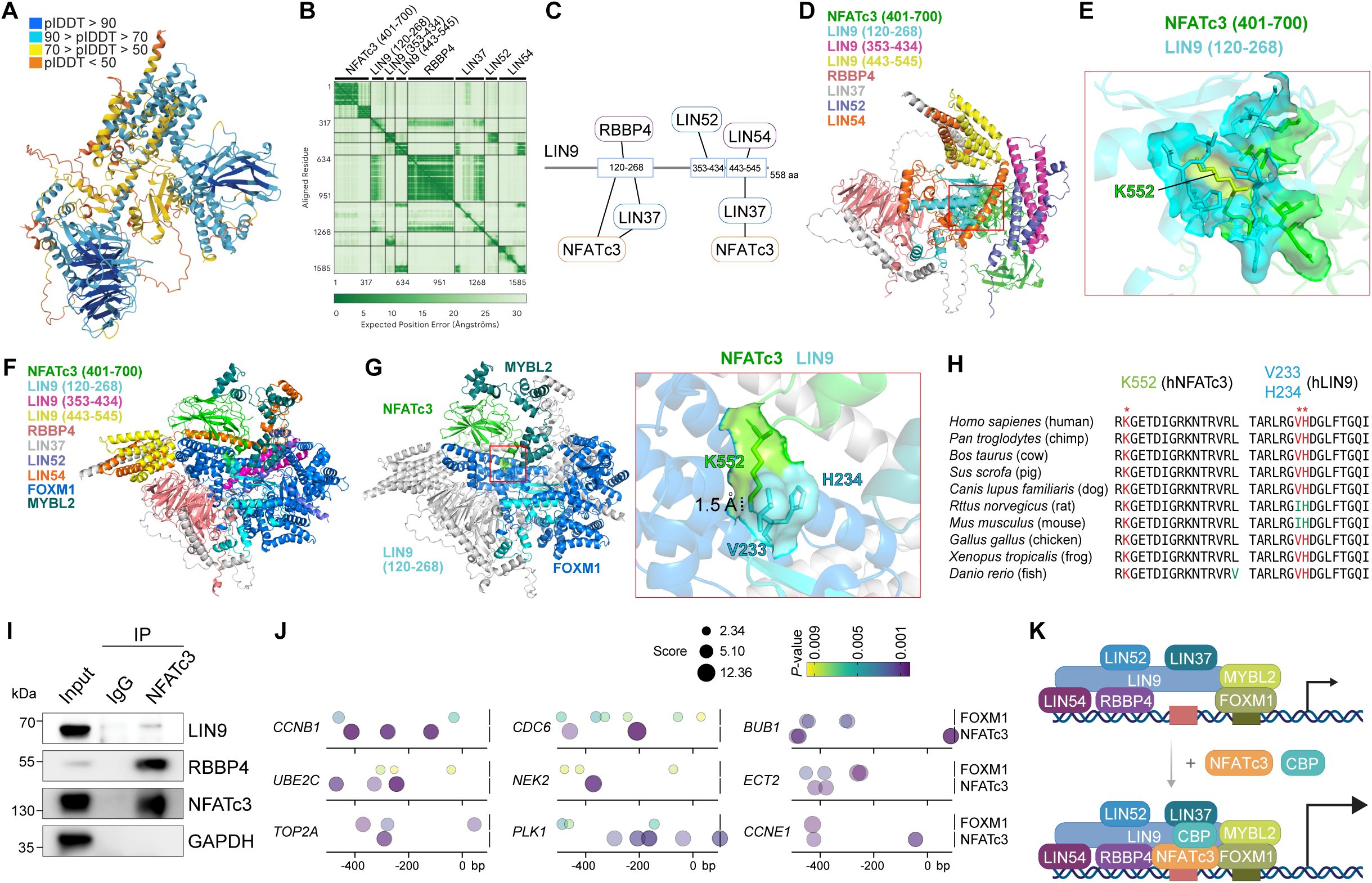
Physical association of NFATc3 with the DREAM complex via the LIN9/LIN37 scaffold. **A, B.** AlphaFold-multimer prediction (A) and corresponding predicted aligned error (PAE) plot (B) of the NFATc3-MuvB complex (LIN9, LIN37, LIN52, LIN54, and RBBP4). Structural confidence is represented by the predicted local distance difference test (plDDT) in (A). **C, D.** Schematic diagram of NFATc3-MuvB complex based on AlphaFold prediction and its visualization using PyMOL. **E**. NFATc3-LIN9 interaction, calculated by protein-protein interaction (PPI) analysis using PyMOL, 3.6 Å distance as a cutoff. **F.** Illustration of NFATc3-MuvB-FoxM1-MYBL2 complex predicted by AlphaFold; visualized by PyMOL. **G.** Interaction interface of NFATc3-FOXM1-LIN9-MYBL2 calculated by PPI analysis. NFATc3 and FOXM1 are highlighted with green and blue, respectively; 1.5 Å: The distance between the residues of K552 (NFATc3) and V233 (LIN9). **H.** Amino acid sequence alignment of NFATc3 and FOXM1 across diverse species**. I.** Co-immunoprecipitation (coIP) in A549 cells using IgG (control) or anti-NFATc3, followed by immunoblot (IB). **J.** Mapping of NFATc3 and FOXM1 binding sites on DREAM target gene promoters. Binding motifs for NFATc3 and FOXM1 were predicted using JASPAR position weight matrices (PWMs) and mapped using FIMO (Find Individual Motif Occurrences) (*P* < 0.01). Bubble size (FIMO score) indicating the match strength between the DNA sequence and the transcription factor’s PWM. Larger bubbles represent stronger consensus match. **K.** Illustration of working model. In the presence of active NFATc3, NFATc3 is recruited to the DREAM target gene promoters via the LIN9/LIN37-based scaffold. CBP is likely to be co-enriched with NFATc3 for enhanced transactivation of the DREAM target genes.

## Discussion

Most genes regulated by the DREAM complex are related to the cell cycle, yet the impact of the DREAM complex on tumorigenesis has not been thoroughly studied. The DREAM complex has recently gained attention for its role in coordinating the cell cycle^46^. We previously found that PCLAF remodels the DREAM complex to activate genes involved in cell hyperproliferation and exit from quiescence in LUAD^17^. Interestingly, our unbiased screening identified two calcium signaling inhibitors that mimic the suppression of DREAM target genes and inhibit LUAD tumorigenesis^17^. These findings prompted us to investigate the oncogenic roles of calcium signaling in LUAD. Indeed, blocking calcium signaling was sufficient to suppress LUAD cell proliferation. Among the five NFATs, NFATc3 was highly expressed in LUAD, and its depletion led to the downregulation of DREAM target genes. Genome-wide analysis further revealed NFATc3 enrichment on the promoters of DREAM target genes, and NFATc3 was shown to interact with DREAM complex components.

Our AlphaFold analysis of the MuvB, a core complex of the DREAM complex, identified that LIN9 serves as a scaffolding protein that binds to LIN37, LIN52, LIN54, and RBBP4 (Fig. 4A-D), which is consistent with the previous study showing the protein structure of the LIN9-LIN37-RBBP4 by X-ray crystallography^45^. Additionally, LIN9 was shown to interact with NFATc3 protein (Fig. 4I). This interaction is particularly significant as it suggests a structural remodeling of the DREAM complex. Typically, the MuvB core (LIN9, LIN37, LIN52, LIN54, and RBBP4) serves as a platform that recruits either the repressive RBL2/p130-E2F4 module during G0/G1 phase or the activating FOXM1/MYBL2 module during S/G2 phase^12, 13^. Our finding that NFATc3 binds to LIN9 indicates that NFATc3 may act as a new regulatory partner that docks onto the MuvB scaffold. This assembly potentially remodels the DREAM complex from its canonical repressive state into a potent transcriptional activator. Previously, we found that harmine, an inhibitor of dual-tyrosine-related kinases (Dyrks) that phosphorylates LIN52 for recruiting the RBL2/p130-E2F4 for the DREAM target gene repression, failed to rescue CsA-induced DREAM target genes repression^17^. This suggests that NFATc3-enhanced gene transactivation of the DREAM complex is somehow independent of the RBL2/p130-E2F4 repressive module. NFATs bind to promoters in conjunction with other transcription factors (TFs), AP-1 and RUNX or CBP/p300 co-activator^47, 48^. Intriguingly, NFATc3 binding sites were located in the proximal promoters of the DREAM target genes (Fig. 4J), implying that NFATc3 and FOXM1 cooperatively transactivate the DREAM target genes. Genome-wide co-occupancy of the DREAM target genes by NFATc3 and FOXM1 needs to be determined. Additionally, key amino acids for NFATc3-LIN9 interaction warrant experimental validation.

NFATs have been extensively studied in immune contexts, where calcium-NFAT signaling plays a crucial role in T cell activation. Consequently, calcium inhibitors have been used for immune suppression in patients undergoing organ transplantation. For instance, CsA is commonly prescribed for organ transplantation or rheumatoid arthritis due to its immune-suppressive effects. Given the role of immune surveillance in targeting tumor cells, the blockade of calcium signaling and subsequent inhibition of T cell activation could act as a double-edged sword. Therefore, long-term blockade of calcium-NFATc3 may not be applicable for LUAD treatment. Instead, short-term CsA use, such as in adjuvant therapy or in combination with other first- or second-line therapies or T cell-based immune checkpoint inhibitors, would be preferable. Furthermore, it would be valuable to investigate how calcium signaling is hyperactivated in LUAD.

Taken together, our study dissects the underlying mechanism of aberrant cell hyperproliferation in LUAD by identifying that calcium-NFATc3 activates the DREAM complex. This finding reveals a new aspect of lung tumorigenesis and proposes the calcium-NFATc3 axis as a molecular target for LUAD with a hyperactivated DREAM pathway.

## Methods

### Cell culture and drugs

Human NSCLC cell lines (A549, H358, H23, H1792, and H1355) were purchased from ATCC. All NSCLC cell lines were grown in RPMI 1640 medium with 10% fetal bovine serum (FBS) and 1% penicillin/streptomycin (10,000 U/mL). A549 and H1792 were treated with 5 µM cyclosporin A (cys; Selleckchem), FK506 (Tacrolimus; Selleckchem), and BAPTA-AM (Selleckchem) for 24 or 48 hours.

### Crystal violet staining

To measure cell proliferation, the same number of cells per well was seeded onto six-well plates in triplicate and grown for 3∼6 days. Cell growth rates were analyzed by daily cell counting with a Bio-Rad TC10 automated cell counter or by measuring optical density (OD values, 590 nm) at the end point after crystal violet staining. For cell growth assays with chemical treatment, fresh chemicals were added to the media every three days. For crystal violet staining, a 0.5% crystal violet solution was prepared by dissolving 0.5 g of crystal violet. Fixed cells were incubated with the crystal violet solution at room temperature for 10 minutes. Excess stain was removed by washing the cells with distilled water 3–4 times until the background was clear.

### Immunostaining

For IHC analysis, paraffin-sectioned samples^17^ were immunostained according to standard protocols (Wang et al., 2016). The following antibodies were used for immunohistochemistry: NAFTc3 (Santa Cruz, sc-8405x). For antigen retrieval, FFPE slices were subjected to heat-induced epitope retrieval pre-treatment at 120°C using citrate-based antigen unmasking solution (Vector Laboratories, Burlingame, CA, USA). For immunofluorescence, after blocking with 10% goat serum in PBS for 30 min at ambient temperature, sections were incubated with primary antibody overnight at 4°C and secondary antibody (1:200) for 1 h at ambient temperature. Sections were mounted with ProLong Gold antifade reagent with DAPI (Invitrogen). For immunohistochemistry, sections were incubated with primary antibody overnight at 4°C and secondary antibodies (1:200) for 1 h at ambient temperature. 3,3’-Diaminobenzidine (DAB) (Vector Laboratory) was used as the chromogen. Then, sections were dehydrated and mounted with Permount (Thermo Fisher Scientific). Images were captured with the fluorescence microscope (Zeiss).

### Plasmids, transfections, and viral infections

Viral infection and selection of human NSCLC cell lines were performed as previously described^29^. Human shRNAs against PCLAF (shPCLAF#1, TRCN0000278496; shPCLAF#2, TRCN0000278497) were used for generating PCLAF KD cells. To obtain stable Pclaf KD murine lung cancer cell lines, shRNAs against Pclaf (GIPZ mouse Pclaf shRNA, Dharmacon; V2LMM_11233, V2LMM_16348) were used. Cell lines transfected with empty vector (GIPZ empty) were used as controls. Plasmids encoding the open reading frames of *PCLAF* and *RBBP4* were obtained from the Functional Genomics Core Facility at MD Anderson Cancer Center. For subcloning, the HA-NFAT4(3-407)-GFP vector (Addgene) was modified using pcDNA-3xFLAG (N terminus) or pLenti-3xFLAG (N terminus)-hygro mammalian expression plasmids, as previously performed^29^. pLenti-3xFLAG (N terminus)-hygro plasmids were used for rescue experiments (hygromycin selection, 150∼200 μg/mL).

### Flow cytometry

For cell cycle analysis, trypsinized cells were fixed with 70% ethanol for 2 h at 20 °C and washed twice with phosphate-buffered saline (PBS). Next, 1×10^6^ cells were resuspended and incubated with RNase A (20 μg/mL) and propidium iodide solution (50 μg/mL) for 30 min. Singlet cells were analyzed by FACS. For cell synchronization assays, cells were arrested in G1/S by double thymidine block and released cells (S phase) were collected at each time point for FACS analysis. For the assessment of cells at G0, freshly ethanol-fixed cells (1×10^6^) were incubated with pyronin Y (1 μg/mL) and 7-aminoactinomycin D (7-AAD; 5 μg/mL) for 45 min at 37°C, and low-RNA-content cells (low pyronin Y signals) were defined as G0 cells on FACS analysis (Schmid et al., 2000). For analysis of the cell cycle phases of NFATc3 KD cells, cells were fixed with 4% paraformaldehyde. Cell cycle phases were analyzed by FACS with 7-AAD and propidium iodide staining. All cells were cultured at 60∼80% confluence.

### qRT-PCR

RNAs were extracted by TRIzol (Invitrogen) and used to synthesize cDNAs using the iScript cDNA synthesis kit (Biorad). qRT-PCR was performed using an Applied Biosystems 7500 Real-Time PCR machine with the primers. Target gene expression was normalized to that of mouse *Hprt1* or human *GAPDH*. Comparative 2^-ΔΔCt^ methods were used for the quantification of qRT-PCR results. The primer sequences are shown below: *PCLAF* (5’-CCAGG GTAAA CAAGG AGACG-3’, 5’-CAGGA AGCAG TGGCT TAGGA-3’), *TOP2A* (5’-GTGAC CCAGC AAATG TGGGT TTACGA-3’, 5’-TGGGT CCCTT TGTTT GTTGT CCGC-3’), *FOXM1* (5’-TCTGA GCGGC CAC CC TACTC TT-3’, 5’-GCCTG GCTTG GCAAT GTGCT-3’), *UBE2C* (5’-CCTGA AGGAA AAGTG GTCTG CCCT-3’, 5’-TCCAG AGCTC GGCAG CATGT-3’), *CCNB1* (5’-ACAGG TCTTC TTCTG CAGGG-3’, 5’-GAACT TGAGC CAGAA CCTGA-3’), *AURKA* (5’-TCCGG CCTCA AACCC AAACC A-3’, 5’-TCTGC TCGCA AAGGG CTCCA-3’), *NEK2* (5’-TCAGT TACAG GAGCG AGAGC GAGC-3’, 5’-CTGCT CTAGC CAGTT TGTCC TCTGC-3’), *ECT2* (5’-CACGG ACTTT CAGGA TTCTG TC-3’, 5’-GAAAA TGGCA AAGGC TCTCC-3’), *BUB1* (5’-AGCAT GCCAG TGCTG TCCTT CA G -3’, 5’-GCAAA TGGGT TTCAG TGAGG CG-3’), *PLK1* (5’-TATCC ATTCA CCGCA GCCTC GC-3’, 5’-TGTGC AGCTC CAGGA GAGAC CT-3’), *KIF4* (5’-GCCTC AGGTG GTGGT TGGTA CA-3’, 5’-CCATA GGCCA GGACC GTTGC AT-3’), *CENPE* (5’-AGCTG GAGAG TTGCA GTTAC TGTTG GA-3’, 5’-CCACC TGGTC TTTGT GCATG CAATCT-3’), *NFATc1* (5’-ACCAG GTGCA CCGCA TCACA-3’, 5’-TCGCA TGCTG TTCTC CGGCA-3’), *NFATc2* (5’-AGGTG CAGCC CAAGC CACAT-3’, 5’-AACCA CAGGG TGGCC TCCAG TT-3’), *NFATc3* (5’-TCACC AGCCC GGGAG ACTTC AA-3’, 5’-GGCCA GGCTT TGGTT TGCTC CA-3’), *NFATc4* (5’-ATCGG TGCCC ATCGA GTGCT-3’, 5’-ACCAGT TCCTC GCCTC CTCTC A-3’), *NFAT5* (5’-GCAGCCGTGG CTCAG TGAAA GA-3’, 5’-TCGTC CAGAG TCGTT GCCCA CA-3’), *GAPDH* (5’-AATGA AGGGG TCATT GATGG-3’, 5’-AAGGT GAAGG TCGGA GTCAA-3’).

### Promoter analysis

BigWig files generated from ChIP-seq were downloaded from the Gene Expression Omnibus (GEO) database and uploaded to the UCSC genome browser (https://genome.ucsc.edu). GEO accession numbers are listed in the figure.

### CUT&RUN analysis

CUTANA ChIC/CUT&RUN Kit (EpiCypher, Cat. No. 14-1048) was used as previously described^49^. In brief, 5’10⁵ cells (A549 cell lines untreated vs. treated with 5 µM CsA for 48 hrs) were pelleted at 600 g for 3 minutes at room temperature (RT). After resuspending the cells twice with 100 µL of washing buffer (pre-wash buffer, protease inhibitors, and 0.5 mM spermidine), the cells were resuspended in wash buffer, preparing them for binding with beads. Next, 100 µL of the cell suspension was added to 10 µL of concanavalin A beads in 8-strip tubes, and the bead-cell slurry was incubated for 10 min at RT. After a brief spin-down, the tubes were placed on a magnet to quickly discard the remaining supernatant. The tubes were then removed from the magnet, and 50 µL of cold antibody buffer (cell permeabilization buffer with 0.01% digitonin and 2 mM EDTA) was immediately added to each reaction. The mixtures were pipetted to resuspend and confirm ConA bead binding. Next, 4 µL of primary antibody (NFATc3 from Santa Cruz) was added to the respective reactions. For the positive and negative control reactions, 1 µL of H3K4me3 positive control antibody and 1 µL of IgG negative control antibody (provided by EpiCypher) were added. Additionally, 2 µL of K-MetStat Panel was added to the reactions designated for the positive and negative control antibodies. The reactions were gently vortexed to mix and incubated overnight on a nutator at 4 °C. After overnight incubation, the tubes were briefly spun, placed on a magnet to allow the slurry to clear, and the supernatant was removed. While keeping the tubes on the magnet, 200 µL of cold cell permeabilization buffer (wash buffer with 0.01% digitonin) was added to each reaction. Next, 2.5 µL of pAG-MNase was added to each reaction, followed by gentle vortexing and a 10 min incubation at RT. The tubes were then quickly spun, and placed on the magnet to clear the slurry, and the supernatant was removed. While keeping the tubes on the magnet, 200 µL of cold cell permeabilization buffer was added directly onto the beads, and the supernatant was removed. The tubes were then removed from the magnet, and 50 µL of cold cell permeabilization buffer was immediately added to each reaction, followed by gentle vortexing to mix and disperse clumps by pipetting. Subsequently, 1 µL of 100 mM calcium chloride was added to each reaction, and the tubes were incubated on a nutator for 2 hours at 4 °C. At the end of the 2-hour incubation, the tubes were quickly spun to collect the liquid, and 34 µL of stop buffer was added to terminate pAG-MNase cleavage activity. The tubes were then placed in a thermocycler set to 37 °C for 10 min. Afterward, the tubes were placed on a magnet, and the supernatants containing CUT&RUN DNA were transferred to new 8-strip tubes. To purify the DNA, 119 µL of SPRIselect beads were slowly added to each reaction, followed by a 5 min incubation at RT. The tubes were then placed on a magnet for 2-5 min at RT, the supernatant was removed, and the beads were washed twice with 180 µL of 85% ethanol. After washing, the tubes were removed from the magnet, and the beads were air-dried for 2-3 min at RT. Finally, 17 µL of 0.1’ TE buffer was added to each reaction to elute the DNA. Then, DNAs were analyzed by qPCR. *ACTB* promoter amplicons served as negative controls. The following primers were used for CUT&RUN PCR: *CCNB1* (5’-CGATC GCCCT GGAAA CGCAT TC-3’, 5’-CCAGC AGAAA CCAAC AGCCG TTC-3’), *TOP2A* (5’-CTCAG CCGTT CATAG GTGGA-3’, 5’-GAACC TTCCT TTAGC CCGCC-3’), *NEK2* (5’-GATCT CGGTT ACCTT GGCGA-3’, 5’-TTAAC CTGTG GGAGA ACCCG-3’), *PLK1* (5’-TGTTC GGGCG TCCGT GTCAA-3’, 5’-ACAAA AGCCT GCGCG CCACT-3’), *UBE2C* (5’-TTGAT TGGTC GACGC CCCCA-3’, 5’-GGTCG CGGTT TTGGG AAGCC AT-3’), *ACTB* (5’-CGGCC AACGC CAAAA CT-3’, 5’-CCCTC TCCCC TCCTT TTGC-3’).

### Co-immunoprecipitation and immunoblotting

For IP, whole-cell lysates were extracted using EBC lysis buffer (50 mM Tris, pH 7.4, 150 mM NaCl, 0.5% NP-40; freshly supplemented with protease and phosphatase inhibitor mixtures) for 30 min at 4°C, followed by centrifugation (12000 g for 10 min). For IP analysis of exogenous protein-protein interactions, whole-cell lysates extracted from stable cells transfected with FLAG-tagged NFATc3 were incubated for 2 h with 15 mL of M2 magnetic beads (Sigma; M8823). For IP analysis of endogenous protein-protein interactions, whole-cell lysates extracted from control or stably transfected H1792 cells were incubated overnight with protein G Dynabeads (Thermo) and 2∼5 ug antibodies against FLAG M2 (Thermo), and NFATc3 (Santa Cruz). After three to five washes with EBC lysis buffer, precipitates were eluted with SDS sample buffer and analyzed using immunoblotting. For analysis of NFATc3 protein levels in NSCLC and NAFTc3 KD cells, radioimmunoprecipitation assay buffer (25mM Tris, pH 7.4, 150mM NaCl, 1% NP-40, 0.5% sodium deoxycholate, 0.1% SDS, and protease and phosphatase inhibitor mixtures) was used for whole-cell lysate preparation.

Immunoblot blocking and antibody incubation were conducted using 2% bovine serum albumin or 5% nonfat dry milk in TBST (25 mM, pH 8.0, 125 mM NaCl, and 0.1% Tween-20). SuperSignal West Pico and Femto (Thermo) were used to detect horseradish peroxidase-conjugated secondary antibodies. The following antibodies were used for immunoblotting: FOXM1 (Santa Cruz, sc-376471x, 1:1000), LIN9 (Santa Cruz, sc-398234, 1:2000), LIN54 (Bethyl, A303-799A-M,1:10000), FLAG M2 (Thermo, F1804, 1:20000) and alpha-tubulin (Cell signaling, 2144S, 1:5000).

### Protein-protein interaction (PPI) analysis

The three-dimensional structures and their potential complex formation were predicted using AlphaFold 3.0^50^. For multimer prediction, the AlphaFold-Multimer pipeline was employed to simulate the interaction interface between NFATc3 (Q12968), LIN9 (Q5TKA1), LIN37 (Q96GY3), LIN52 (Q52LA3), LIN54 (Q6MZP7), RBBP4 (Q09028), FOXM1 (O08696), and MYBL2 (P10244). The amino acid sequences were obtained from the UniProt database. The prediction was performed using five models, and the highest-confidence model, determined by the predicted local distance difference test (plDDT) and predicted aligned error (PAE) scores, was selected for further structural analysis. The predicted complex structure (mmCIF format) was imported into PyMOL 3.1.6.2 for visualization and detailed PPI analysis. Residues at the interface were identified using a distance cutoff (3.6 Å). The final high-resolution images for structural comparison and interaction site mapping were rendered using PyMOL’s ray-tracing feature.

## Author contributions

J.A.: Methodology, investigation, visualization, writing (original draft); Y.S.: Methodology, investigation, visualization, writing (original draft); J.J.: Methodology, investigation, visualization, software, analysis, data curation, writing (original draft); M.J.K.: methodology, investigation; B.K.: Investigation; J.J.: Investigation; S.J.: Investigation; J.-I.P.: Conceptualization, methodology, investigation, analysis, writing (original draft, review, and editing), visualization, supervision, project administration, funding acquisition

## Acknowledgments

This work was supported by the National Cancer Institute (R03 CA279867).

## Supplementary Information

**Supplementary Figure 1.**
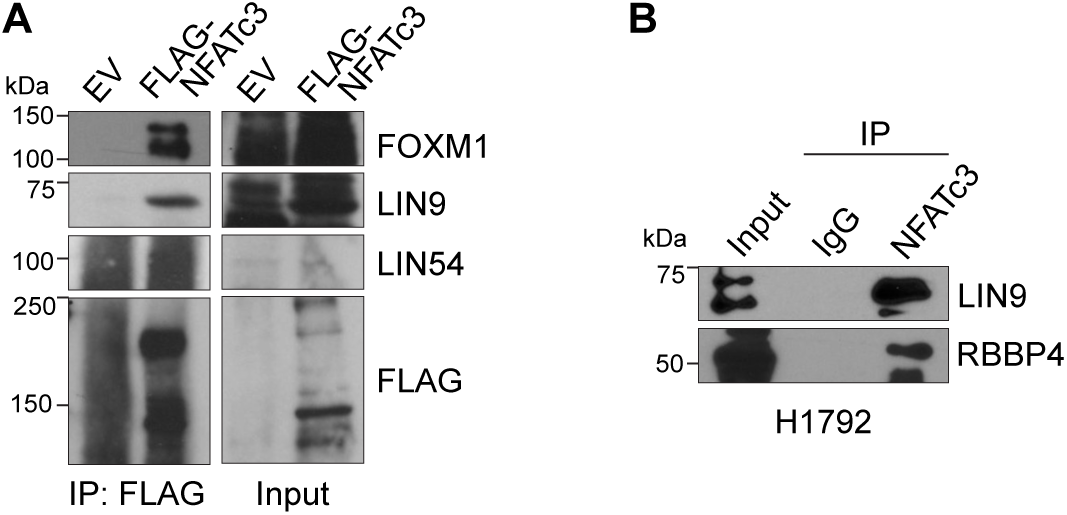
Physical interaction of NFATc3 with the MuvB complex and FOXM1. **A.** Ectopic expression and co-immunoprecipitation (coIP) in H1792 cells. Cells were transiently transfected with empty vector, control FLAG, or FLAG-tagged NFATc3. Cell lysates were immunoprecipitated with an anti-FLAG, followed by immunoblotting (IB) with antibodies against FOXM1, LIN9, and LIN54. **B.** Endogenous co-immunoprecipitation (coIP) in H1792 cells using IgG (control) or anti-NFATc3. Non-specific IgG served as a negative control.

## References

1. Travis WD, Brambilla E, Nicholson AG, Yatabe Y, Austin JHM, Beasley MB, Chirieac LR, Dacic S, Duhig E, Flieder DB, Geisinger K, Hirsch FR, Ishikawa Y, Kerr KM, Noguchi M, Pelosi G, Powell CA, Tsao MS, Wistuba I, Panel WHO. The 2015 World Health Organization Classification of Lung Tumors: Impact of Genetic, Clinical and Radiologic Advances Since the 2004 Classification. J Thorac Oncol. 2015;10(9):1243–60. Epub 2015/08/21. doi: 10.1097/JTO.0000000000000630. PubMed PMID: 26291008.

2. Cancer Genome Atlas Research N. Comprehensive molecular profiling of lung adenocarcinoma. Nature. 2014;511(7511):543–50. Epub 2014/08/01. doi: 10.1038/nature13385. PubMed PMID: 25079552; PMCID: PMC4231481.

3. Jackson EL, Willis N, Mercer K, Bronson RT, Crowley D, Montoya R, Jacks T, Tuveson DA. Analysis of lung tumor initiation and progression using conditional expression of oncogenic K-ras. Genes Dev. 2001;15(24):3243–8. Epub 2001/12/26. doi: 10.1101/gad.943001. PubMed PMID: 11751630; PMCID: PMC312845.

4. Jackson EL, Olive KP, Tuveson DA, Bronson R, Crowley D, Brown M, Jacks T. The differential effects of mutant p53 alleles on advanced murine lung cancer. Cancer Res. 2005;65(22):10280–8. Epub 2005/11/17. doi: 10.1158/0008-5472.CAN-05-2193. PubMed PMID: 16288016.

5. Cox AD, Fesik SW, Kimmelman AC, Luo J, Der CJ. Drugging the undruggable RAS: Mission possible? Nat Rev Drug Discov. 2014;13(11):828–51. Epub 2014/10/18. doi: 10.1038/nrd4389. PubMed PMID: 25323927; PMCID: PMC4355017.

6. Muller PA, Vousden KH. Mutant p53 in cancer: new functions and therapeutic opportunities. Cancer Cell. 2014;25(3):304–17. Epub 2014/03/22. doi: 10.1016/j.ccr.2014.01.021. PubMed PMID: 24651012; PMCID: PMC3970583.

7. Malumbres M, Barbacid M. To cycle or not to cycle: a critical decision in cancer. Nat Rev Cancer. 2001;1(3):222–31. Epub 2002/03/21. doi: 10.1038/35106065. PubMed PMID: 11902577.

8. Hanahan D, Weinberg RA. Hallmarks of cancer: the next generation. Cell. 2011;144(5):646–74. Epub 2011/03/08. doi: 10.1016/j.cell.2011.02.013. PubMed PMID: 21376230.

9. Lewis PW, Beall EL, Fleischer TC, Georlette D, Link AJ, Botchan MR. Identification of a Drosophila Myb-E2F2/RBF transcriptional repressor complex. Genes Dev. 2004;18(23):2929–40. Epub 2004/11/17. doi: 10.1101/gad.1255204. PubMed PMID: 15545624; PMCID: PMC534653.

10. Korenjak M, Taylor-Harding B, Binne UK, Satterlee JS, Stevaux O, Aasland R, White-Cooper H, Dyson N, Brehm A. Native E2F/RBF complexes contain Myb-interacting proteins and repress transcription of developmentally controlled E2F target genes. Cell. 2004;119(2):181–93. Epub 2004/10/14. doi: 10.1016/j.cell.2004.09.034. PubMed PMID: 15479636.

11. Harrison MM, Ceol CJ, Lu X, Horvitz HR. Some C. elegans class B synthetic multivulva proteins encode a conserved LIN-35 Rb-containing complex distinct from a NuRD-like complex. Proc Natl Acad Sci U S A. 2006;103(45):16782–7. Epub 2006/11/01. doi: 10.1073/pnas.0608461103. PubMed PMID: 17075059; PMCID: PMC1636532.

12. Litovchick L, Sadasivam S, Florens L, Zhu X, Swanson SK, Velmurugan S, Chen R, Washburn MP, Liu XS, DeCaprio JA. Evolutionarily conserved multisubunit RBL2/p130 and E2F4 protein complex represses human cell cycle-dependent genes in quiescence. Mol Cell. 2007;26(4):539–51. Epub 2007/05/29. doi: 10.1016/j.molcel.2007.04.015. PubMed PMID: 17531812.

13. Sadasivam S, Duan S, DeCaprio JA. The MuvB complex sequentially recruits B-Myb and FoxM1 to promote mitotic gene expression. Genes Dev. 2012;26(5):474–89. Epub 2012/03/07. doi: 10.1101/gad.181933.111. PubMed PMID: 22391450; PMCID: PMC3305985.

14. Sadasivam S, DeCaprio JA. The DREAM complex: master coordinator of cell cycle-dependent gene expression. Nat Rev Cancer. 2013;13(8):585–95. Epub 2013/07/12. doi: 10.1038/nrc3556. PubMed PMID: 23842645; PMCID: PMC3986830.

15. MacDonald J, Ramos-Valdes Y, Perampalam P, Litovchick L, DiMattia GE, Dick FA. A Systematic Analysis of Negative Growth Control Implicates the DREAM Complex in Cancer Cell Dormancy. Mol Cancer Res. 2017;15(4):371–81. Epub 2016/12/30. doi: 10.1158/1541-7786.MCR-16-0323-T. PubMed PMID: 28031411.

16. Nor Rashid N, Yusof R, Watson RJ. Disruption of repressive p130-DREAM complexes by human papillomavirus 16 E6/E7 oncoproteins is required for cell-cycle progression in cervical cancer cells. J Gen Virol. 2011;92(Pt 11):2620–7. Epub 2011/08/05. doi: 10.1099/vir.0.035352-0. PubMed PMID: 21813705.

17. Kim MJ, Cervantes C, Jung YS, Zhang X, Zhang J, Lee SH, Jun S, Litovchick L, Wang W, Chen J, Fang B, Park JI. PAF remodels the DREAM complex to bypass cell quiescence and promote lung tumorigenesis. Mol Cell. 2021;81(8):1698–714 e6. Epub 20210223. doi: 10.1016/j.molcel.2021.02.001. PubMed PMID: 33626321; PMCID: PMC8052288.

18. Yu P, Huang B, Shen M, Lau C, Chan E, Michel J, Xiong Y, Payan DG, Luo Y. p15(PAF), a novel PCNA associated factor with increased expression in tumor tissues. Oncogene. 2001;20(4):484–9. Epub 2001/04/21. doi: 10.1038/sj.onc.1204113. PubMed PMID: 11313979.

19. Emanuele MJ, Ciccia A, Elia AE, Elledge SJ. Proliferating cell nuclear antigen (PCNA)-associated KIAA0101/PAF15 protein is a cell cycle-regulated anaphase-promoting complex/cyclosome substrate. Proc Natl Acad Sci U S A. 2011;108(24):9845–50. Epub 2011/06/02. doi: 10.1073/pnas.1106136108. PubMed PMID: 21628590; PMCID: PMC3116415.

20. Povlsen LK, Beli P, Wagner SA, Poulsen SL, Sylvestersen KB, Poulsen JW, Nielsen ML, Bekker-Jensen S, Mailand N, Choudhary C. Systems-wide analysis of ubiquitylation dynamics reveals a key role for PAF15 ubiquitylation in DNA-damage bypass. Nat Cell Biol. 2012;14(10):1089–98. Epub 2012/09/25. doi: 10.1038/ncb2579. PubMed PMID: 23000965.

21. Cheng Y, Li K, Diao D, Zhu K, Shi L, Zhang H, Yuan D, Guo Q, Wu X, Liu D, Dang C. Expression of KIAA0101 protein is associated with poor survival of esophageal cancer patients and resistance to cisplatin treatment in vitro. Lab Invest. 2013;93(12):1276–87. Epub 2013/10/23. doi: 10.1038/labinvest.2013.124. PubMed PMID: 24145239.

22. Hosokawa M, Takehara A, Matsuda K, Eguchi H, Ohigashi H, Ishikawa O, Shinomura Y, Imai K, Nakamura Y, Nakagawa H. Oncogenic role of KIAA0101 interacting with proliferating cell nuclear antigen in pancreatic cancer. Cancer Res. 2007;67(6):2568–76. Epub 2007/03/17. doi: 10.1158/0008-5472.CAN-06-4356. PubMed PMID: 17363575.

23. Jain M, Zhang L, Patterson EE, Kebebew E. KIAA0101 is overexpressed, and promotes growth and invasion in adrenal cancer. PLoS One. 2011;6(11):e26866. Epub 2011/11/19. doi: 10.1371/journal.pone.0026866. PubMed PMID: 22096502; PMCID: PMC3214018.

24. Yuan RH, Jeng YM, Pan HW, Hu FC, Lai PL, Lee PH, Hsu HC. Overexpression of KIAA0101 predicts high stage, early tumor recurrence, and poor prognosis of hepatocellular carcinoma. Clin Cancer Res. 2007;13(18 Pt 1):5368–76. Epub 2007/09/19. doi: 10.1158/1078-0432.CCR-07-1113. PubMed PMID: 17875765.

25. Mizutani K, Onda M, Asaka S, Akaishi J, Miyamoto S, Yoshida A, Nagahama M, Ito K, Emi M. Overexpressed in anaplastic thyroid carcinoma-1 (OEATC-1) as a novel gene responsible for anaplastic thyroid carcinoma. Cancer. 2005;103(9):1785–90. Epub 2005/03/25. doi: 10.1002/cncr.20988. PubMed PMID: 15789362.

26. Wang X, Jung YS, Jun S, Lee S, Wang W, Schneider A, Sun Oh Y, Lin SH, Park BJ, Chen J, Keyomarsi K, Park JI. PAF-Wnt signaling-induced cell plasticity is required for maintenance of breast cancer cell stemness. Nat Commun. 2016;7:10633. Epub 20160204. doi: 10.1038/ncomms10633. PubMed PMID: 26843124; PMCID: PMC4743006.

27. Yan R, Zhu K, Dang C, Lan K, Wang H, Yuan D, Chen W, Meltzer SJ, Li K. Paf15 expression correlates with rectal cancer prognosis, cell proliferation and radiation response. Oncotarget. 2016;7(25):38750–61. Epub 2016/10/23. doi: 10.18632/oncotarget.9606. PubMed PMID: 27246972; PMCID: PMC5122426.

28. Jun S, Lee S, Kim HC, Ng C, Schneider AM, Ji H, Ying H, Wang H, DePinho RA, Park JI. PAF-mediated MAPK signaling hyperactivation via LAMTOR3 induces pancreatic tumorigenesis. Cell Rep. 2013;5(2):314–22. doi: 10.1016/j.celrep.2013.09.026. PubMed PMID: 24209743; PMCID: PMC4157353.

29. Jung HY, Jun S, Lee M, Kim HC, Wang X, Ji H, McCrea PD, Park JI. PAF and EZH2 induce Wnt/beta-catenin signaling hyperactivation. Mol Cell. 2013;52(2):193–205. Epub 20130919. doi: 10.1016/j.molcel.2013.08.028. PubMed PMID: 24055345; PMCID: PMC4040269.

30. Kim MJ, Xia B, Suh HN, Lee SH, Jun S, Lien EM, Zhang J, Chen K, Park JI. PAF-Myc-Controlled Cell Stemness Is Required for Intestinal Regeneration and Tumorigenesis. Dev Cell. 2018;44(5):582–96 e4. doi: 10.1016/j.devcel.2018.02.010. PubMed PMID: 29533773; PMCID: PMC5854208.

31. Ong DST, Hu B, Ho YW, Sauve CG, Bristow CA, Wang Q, Multani AS, Chen P, Nezi L, Jiang S, Gorman CE, Monasterio MM, Koul D, Marchesini M, Colla S, Jin EJ, Sulman EP, Spring DJ, Yung WA, Verhaak RGW, Chin L, Wang YA, DePinho RA. PAF promotes stemness and radioresistance of glioma stem cells. Proc Natl Acad Sci U S A. 2017;114(43):E9086–E95. Epub 2017/10/27. doi: 10.1073/pnas.1708122114. PubMed PMID: 29073105; PMCID: PMC5664518.

32. Kim B, Huang Y, Ko KP, Zhang S, Zou G, Zhang J, Kim MJ, Little D, Ellis LV, Paschini M, Jun S, Park KS, Chen J, Kim C, Park JI. PCLAF-DREAM drives alveolar cell plasticity for lung regeneration. Nat Commun. 2024;15(1):9169. Epub 20241024. doi: 10.1038/s41467-024-53330-1. PubMed PMID: 39448571; PMCID: PMC11502753.

33. Subramanian A, Narayan R, Corsello SM, Peck DD, Natoli TE, Lu X, Gould J, Davis JF, Tubelli AA, Asiedu JK, Lahr DL, Hirschman JE, Liu Z, Donahue M, Julian B, Khan M, Wadden D, Smith IC, Lam D, Liberzon A, Toder C, Bagul M, Orzechowski M, Enache OM, Piccioni F, Johnson SA, Lyons NJ, Berger AH, Shamji AF, Brooks AN, Vrcic A, Flynn C, Rosains J, Takeda DY, Hu R, Davison D, Lamb J, Ardlie K, Hogstrom L, Greenside P, Gray NS, Clemons PA, Silver S, Wu X, Zhao WN, Read-Button W, Wu X, Haggarty SJ, Ronco LV, Boehm JS, Schreiber SL, Doench JG, Bittker JA, Root DE, Wong B, Golub TR. A Next Generation Connectivity Map: L1000 Platform and the First 1,000,000 Profiles. Cell. 2017;171(6):1437–52 e17. Epub 2017/12/02. doi: 10.1016/j.cell.2017.10.049. PubMed PMID: 29195078; PMCID: PMC5990023.

34. Clapham DE. Calcium signaling. Cell. 2007;131(6):1047–58. Epub 2007/12/18. doi: 10.1016/j.cell.2007.11.028. PubMed PMID: 18083096.

35. Huang L, Keyser BM, Tagmose TM, Hansen JB, Taylor JT, Zhuang H, Zhang M, Ragsdale DS, Li M. NNC 55-0396 [(1S,2S)-2-(2-(N-[(3-benzimidazol-2-yl)propyl]-N-methylamino)ethyl)-6-fluoro-1,2, 3,4-tetrahydro-1-isopropyl-2-naphtyl cyclopropanecarboxylate dihydrochloride]: a new selective inhibitor of T-type calcium channels. J Pharmacol Exp Ther. 2004;309(1):193–9. Epub 2004/01/14. doi: 10.1124/jpet.103.060814. PubMed PMID: 14718587.

36. Kim KH, Kim D, Park JY, Jung HJ, Cho YH, Kim HK, Han J, Choi KY, Kwon HJ. NNC 55-0396, a T-type Ca2+ channel inhibitor, inhibits angiogenesis via suppression of hypoxia-inducible factor-1alpha signal transduction. J Mol Med (Berl). 2015;93(5):499–509. Epub 2014/12/05. doi: 10.1007/s00109-014-1235-1. PubMed PMID: 25471482.

37. Dziegielewska B, Brautigan DL, Larner JM, Dziegielewski J. T-type Ca2+ channel inhibition induces p53-dependent cell growth arrest and apoptosis through activation of p38-MAPK in colon cancer cells. Mol Cancer Res. 2014;12(3):348–58. Epub 2013/12/24. doi: 10.1158/1541-7786.MCR-13-0485. PubMed PMID: 24362252.

38. Huang W, Lu C, Wu Y, Ouyang S, Chen Y. T-type calcium channel antagonists, mibefradil and NNC-55-0396 inhibit cell proliferation and induce cell apoptosis in leukemia cell lines. J Exp Clin Cancer Res. 2015;34:54. Epub 2015/05/21. doi: 10.1186/s13046-015-0171-4. PubMed PMID: 25989794; PMCID: PMC4443536.

39. Handschumacher RE, Harding MW, Rice J, Drugge RJ, Speicher DW. Cyclophilin: a specific cytosolic binding protein for cyclosporin A. Science. 1984;226(4674):544–7. Epub 1984/11/02. doi: 10.1126/science.6238408. PubMed PMID: 6238408.

40. Stewart TA, Yapa KT, Monteith GR. Altered calcium signaling in cancer cells. Biochim Biophys Acta. 2015;1848(10 Pt B):2502–11. Epub 2014/08/26. doi: 10.1016/j.bbamem.2014.08.016. PubMed PMID: 25150047.

41. Monteith GR, Davis FM, Roberts-Thomson SJ. Calcium channels and pumps in cancer: changes and consequences. J Biol Chem. 2012;287(38):31666–73. Epub 2012/07/24. doi: 10.1074/jbc.R112.343061. PubMed PMID: 22822055; PMCID: PMC3442501.

42. Sallan MC, Visa A, Shaikh S, Nager M, Herreros J, Canti C. T-type Ca(2+) Channels: T for Targetable. Cancer Res. 2018;78(3):603–9. Epub 2018/01/19. doi: 10.1158/0008-5472.CAN-17-3061. PubMed PMID: 29343521.

43. Lin Y, Song Y, Zhang Y, Shi M, Hou A, Han S. NFAT signaling dysregulation in cancer: Emerging roles in cancer stem cells. Biomed Pharmacother. 2023;165:115167. Epub 20230714. doi: 10.1016/j.biopha.2023.115167. PubMed PMID: 37454598.

44. Zheng S, Wang X, Zhao D, Liu H, Hu Y. Calcium homeostasis and cancer: insights from endoplasmic reticulum-centered organelle communications. Trends Cell Biol. 2023;33(4):312–23. Epub 20220729. doi: 10.1016/j.tcb.2022.07.004. PubMed PMID: 35915027.

45. Asthana A, Ramanan P, Hirschi A, Guiley KZ, Wijeratne TU, Shelansky R, Doody MJ, Narasimhan H, Boeger H, Tripathi S, Muller GA, Rubin SM. The MuvB complex binds and stabilizes nucleosomes downstream of the transcription start site of cell-cycle dependent genes. Nat Commun. 2022;13(1):526. Epub 20220126. doi: 10.1038/s41467-022-28094-1. PubMed PMID: 35082292; PMCID: PMC8792015.

46. Walston H, Iness AN, Litovchick L. DREAM On: Cell Cycle Control in Development and Disease. Annu Rev Genet. 2021;55:309–29. Epub 20210908. doi: 10.1146/annurev-genet-071819-103836. PubMed PMID: 34496610.

47. Gabriel CH, Gross F, Karl M, Stephanowitz H, Hennig AF, Weber M, Gryzik S, Bachmann I, Hecklau K, Wienands J, Schuchhardt J, Herzel H, Radbruch A, Krause E, Baumgrass R. Identification of Novel Nuclear Factor of Activated T Cell (NFAT)-associated Proteins in T Cells. J Biol Chem. 2016;291(46):24172–87. Epub 2016/09/18. doi: 10.1074/jbc.M116.739326. PubMed PMID: 27637333; PMCID: PMC5104941.

48. Lee JU, Kim LK, Choi JM. Revisiting the Concept of Targeting NFAT to Control T Cell Immunity and Autoimmune Diseases. Front Immunol. 2018;9:2747. Epub 2018/12/13. doi: 10.3389/fimmu.2018.02747. PubMed PMID: 30538703; PMCID: PMC6277705.

49. Seo Y, Zhang S, Jang J, Ko KP, Kim KB, Huang Y, Kim DW, Kim B, Zou G, Zhang J, Jun S, Chu W, Kirk NA, Hwang YE, Ban YH, Dhar SS, Chan JM, Lee MG, Rudin CM, Park KS, Park JI. Actin dysregulation induces neuroendocrine plasticity and immune evasion: a vulnerability of small cell lung cancer. Nat Commun. 2025. Epub 20251206. doi: 10.1038/s41467-025-67078-9. PubMed PMID: 41353272.

50. Abramson J, Adler J, Dunger J, Evans R, Green T, Pritzel A, Ronneberger O, Willmore L, Ballard AJ, Bambrick J, Bodenstein SW, Evans DA, Hung CC, O’Neill M, Reiman D, Tunyasuvunakool K, Wu Z, Zemgulyte A, Arvaniti E, Beattie C, Bertolli O, Bridgland A, Cherepanov A, Congreve M, Cowen-Rivers AI, Cowie A, Figurnov M, Fuchs FB, Gladman H, Jain R, Khan YA, Low CMR, Perlin K, Potapenko A, Savy P, Singh S, Stecula A, Thillaisundaram A, Tong C, Yakneen S, Zhong ED, Zielinski M, Zidek A, Bapst V, Kohli P, Jaderberg M, Hassabis D, Jumper JM. Accurate structure prediction of biomolecular interactions with AlphaFold 3. Nature. 2024;630(8016):493–500. Epub 20240508. doi: 10.1038/s41586-024-07487-w. PubMed PMID: 38718835; PMCID: PMC11168924.

